# Transcriptional profiling elucidates the essential role of glycogen synthase kinase 3 to fruiting body formation in *Coprinopsis cinerea*

**DOI:** 10.1101/492397

**Authors:** Kathy PoLam Chan, JinHui Chang, YiChun Xie, Man Kit Cheung, Ka Lee Ma, Hoi Shan Kwan

## Abstract

The functions of glycogen synthase kinase 3 (GSK3) have been well-studied in animal, plant and yeast. However, information on its roles in basidiomycetous fungi is still limited. In this study, we used the model mushroom *Coprinopsis cinerea* to study the characteristics of GSK3 in fruiting body development. Application of a GSK3 inhibitor Lithium chloride (LiCl) induced enhanced mycelial growth and inhibited fruiting body formation in *C. cinerea*. RNA-Seq of LiCl-treated *C. cinerea* resulted in a total of 14128 unigenes. There were 1210 differentially expressed genes (DEGs) between the LiCl-treated samples and control samples in the mycelium stage (first time point), whereas 1402 DEGs were detected at the stage when the control samples formed hyphal knots and the treatment samples were still in mycelium (second time point). Kyoto Encyclopedia of Genes and Genome (KEGG) pathway enrichment analysis of the DEGs revealed significant associations between the enhanced mycelium growth in LiCl treated *C. cinerea* and metabolism pathways such as “biosynthesis of secondary metabolite” and “biosynthesis of antibiotics”. In addition, DEGs involved in cellular process pathways, including “cell cycle-yeast” and “meiosis-yeast”, were identified in *C. cinerea* fruiting body formation suppressed by LiCl under favorable environmental conditions. Our findings suggest that GSK3 activity is essential for fruiting body formation as it affects the expression of fruiting body induction genes and genes in cellular processes. Further functional studies of GSK3 in basidiomycetous fungi may help understand the relationships between environmental signals and fruiting body development.

## Introduction

*Coprinopsis cinerea* is a model organism for studying the development and growth regulation in homobasidiomycetous fungi. It can complete its life cycle in only two weeks under laboratory conditions. There are at least six developmental stages in the life cycle of *C. cinerea:* mycelium, initial, stage 1 primordium, stage 2 primordium, young fruiting body, and mature fruiting body with spore formation. Its gill and frequently the entire cap autolyzes to have an inky black fluid at maturity [1]. Due to its rapid morphogenesis, *C. cinerea* is a favorable model to be used in genetic studies of fruiting body development [2]. In particular, strain #326 (AmutBmut pab1-1) is a homokaryotic strain that forms fruiting body without mating, making it especially suitable for molecular studies of basidomycetous fungi under different environmental conditions [3].

The development of *C. cinerea* depends on the sensing of environmental conditions, including light, temperature, humidity, and nutrients [1,37]. After the mycelium reaches the surface of horse dung, the habitat changes drastically, especially the day-night change of light, temperature and humidity. Circadian rhythm, especially light, not only serves as a trigger for adaptation to this changing environment, but also regulates the sexual development. The transformation from mycelium to hyphal knot requires light, glucose depletion and lower temperature [1]. Blue light is required to enter fruiting body initiation, primordial maturation and karyogamy, which ensures spores to be eaten and spread by animals in the early morning [1,4]. How environmental signals stimulate regulators of adaptation and reproduction has attracted wide research interests. Kinases and transcription factors are among the most intensively investigated [37,38].

Glycogen synthase kinase, GSK3, was initially identified as a serine/threonine protein kinase of the CMGC family that phosphorylates and inactivates glycogen synthase [5]. Studies of GSK3 in mammalian cells have indicated its involvement in many signaling pathways, including cell proliferation and differentiation, transcription factor and microtubule dynamics. As a result, GSK3 is now considered as a central regulator among signaling pathways [6]. Apart from in mammals, GSK3 also has a wide variety of essential cell signaling role in plant and fungi. In plant, GSK3 integrates multiple hormonal signals, such as in hormone crosstalk, to regulate plant stress tolerance and acts as an important component to suppress xylem in the TDIF-TDR signaling pathway [7,8]. GSK3 also regulates the growth, conidiogenesis, mycotoxin production, pathogenicity, and stress response mechanism in phytopathogenic fungi, such as *Fusarium graminearum* and *Magnaporthe oryzae* [9,10], and is considered a control regulator in cellular processes. Moreover, genes encoded as glycogen synthase kinase in ascomycetous fungi, such as *Saccharomyces cerevisiae* and *Neurospora crassa*, regulate protein-protein interaction, meiotic gene activation, and transcription factors of circadian clock under environmental stress like nutrient, temperature and light [11,12,13]. Until now, there is little knowledge on the role of GSK3 in mushroom-forming fungi. Lithium chloride (LiCl), a mood stabilizer for mental illnesses treatment, has been used to study the characteristics of glycogen synthase kinase in human and other animal models [14,15]. The inhibition of GSK3 is the main action of LiCl from pharmacological and genetic studies [16]. LiCl inactivates the phosphorylation of GSK3 to its downstream targets. In this study, we used high-throughput RNA sequencing (RNA-Seq) to reveal the transcriptional response of *C. cinerea* on fruiting body formation and development with GSK3 inhibited by the addition of LiCl. The transcriptional profile revealed could elucidate the role of GSK3 activity in fruiting body development.

## Material and methods

### Strain and cultivation conditions

*C. cinerea* strain #326 (A43mut B43mut pab1-1), a homokaryotic fruiting strain, was grown on yeast extract-malt extract-glucose (YMG) medium solidified with 1.5% (w/v) agar [17] at 37°C for 5-6 days until reaching the edge of plates. In the treatment experiments, a piece of mycelium 5 mm diameter punch from the stock plates was inoculated on plates spread with different concentrations (0.25 μM, 0.5 μM, 1 μM, 3 μM, 5 μM) of LiCl (Sigma-Aldrich) after filter sterilization and covered with cellophane. Autoclaved water was added in the controls. Mycelium in the treatment was incubated at 37°C in total darkness for 4 days and then harvested for RNA extraction and transcriptome sequencing. Another set of RNA extraction was carried out from the control plates after the formation of hyphal knots. After the mycelium reached the edge of plates, the plates were incubated at 28°C with the 12/12 h light/dark cycle for fruiting body development. Each treatment was in triplicate under the same experiment conditions.

### RNA isolation, library preparation and sequencing

For transcriptome sequencing, samples in two time points of *C. cinerea* were harvested, first one in the mycelium stage were harvested after 4 days of incubation in total darkness, before reaching the edge of plates, whereas second one in the hyphal knot stage were collected after 3-4 days of incubation under the 12/12 h light/dark cycle after the mycelium reached the edge of plates. Samples collected were frozen in liquid nitrogen and stored at −80°C before RNA extraction by using the RNeasy Plant Mini Kit (Qiagen). RNA was first treated with the TURBO DNA-free kit according to the manufacturer’s instructions. Quality and quantity of the extracted RNA were assessed on 1.5% agarose gels and an Agilent 2100 bioanalyzer with the Agilent RNA 6000 Nano Kit. RNA with a RNA integrity number (RIN) greater than 8.0 was used for transcriptome library construction. RNA sequencing libraries were constructed by using the TruSeq RNA Sample Prep Kit v2 (Illumina) according to the manufacturer’s instructions. Qualified libraries were amplified on cBot to generate clusters on the flowcell (TruSeq PE Cluster Kit V3–cBot–HS, Illumina). The amplified flowcell was then sequenced paired-end on the HiSeq 2000 System (TruSeq SBS KIT-HS V3, Illumina). The entire process was commissioned and performed by the Beijing Genomics Institute (BGI). The RNA-seq row data was submitted to Gene Expression Omnibus (GEO) and its accession number will be provided later.

### Reads alignment and transcript assembly

Fastp was used to filter out low quality and unclean reads [18]. Hisat2 was used to align the high quality and clear reads against the reference genome from JGI (https://genome.jgi.doe.gov/programs/fungi/index.jsf) [3,19]. All transcripts were defined as unigenes and annotated using the Gene Ontology (GO) database for defining their roles in biological processes, molecular functions and cellular components. The unigenes were also annotated using the EuKaryotic Orthologous Groups (KOG) database to define their roles in metabolism, cellular processes and signaling, information storage and processing.

### Differential expression analysis and gene functional annotation

After mapping to the reference genome, the gene expression levels of each sample were calculated by Cufflinks [20]. Differentially expressed genes (DEGs) between the treatment and control samples were then calculated using the Cuffdiff function in Cufflinks [21]. Only transcripts with a p-value <0.05 and fold change >1 or <-1 were considered statistically significant and used in GO and Kyoto Encyclopedia of Genes and Genome (KEGG) enrichment analyses.

### GO and KEGG enrichment analyses

GO and KEGG enrichment of DEGs were performed using the R package clusterProfiler [22]. The enrichGo function in clusterProfiler was used to identify the hypergeometric distribution of DEGs based on the org.Cci.sgd.db Bioconductor annotation package [22], whereas the enrichKEGG function was used to calculate the statistical enrichment of DEGs for KEGG pathways. The threshold of p-value was set at 0.05 and the p.adjust method of Benjamini & Hochberg (BH) was used to control the false discovery rate [39].

### cDNA synthesis and qRT-PCR validation

Quantitative real time PCR (qRT-PCR) of selected genes from enriched KEGG pathways was used to validate the RNA-Seq data. Approximately 500 ng of total RNA was used to synthesize cDNA using the iScript gDNA Clear cDNA Synthesis Kit (Bio-Rad) according to the manufacturer’s instructions. SYBR green–based qRT-PCR was then performed using SsoAdvanced Universal SYBR Green Supermix (Bio-Rad) on Applied Biosystems 7500. Beta-tubulin was used as an endogenous control for normalization. Detail information of the primers used is in S1 Table.

### Statistical analysis

Data are presented as mean ± SD of three replicates for each treatment. Statistical analysis was carried out using one-way analysis of variance (ANOVA) with p-values < 0.05 considered significant.

## Result

### Effect of LiCl on mycelium growth

Different concentrations of LiCl (0.25 μM, 0.5 μM, 1 μM, 3 μM and 5 μM; water in control) were spread on the surface of YMG plates, on which a block of mycelium from the stock plates was inoculated. After 24 hours, no difference in the colony size was observed between the plates (Fig. 1). However, after 48 hours, *C. cinerea* mycelium treated with 1 μM and 3 μM LiCl grew larger colonies than that in other concentrations of LiCl and control plates. 2-3 days later, the mycelium treated with 0.25 μM, 0.5 μM and 5 μM LiCl grew larger colonies than that of the control. Consecutively, the mycelium treated with 1 μM and 3 μM LiCl reached the edge of plates after 5 days of incubation. However, other plates required 6 days to reach the edge of plates. This suggests that the mycelium in 1 μM and 3 μM LiCl plates had higher growth rate than that in other concentrations of LiCl and control plates.

**Fig.1.**
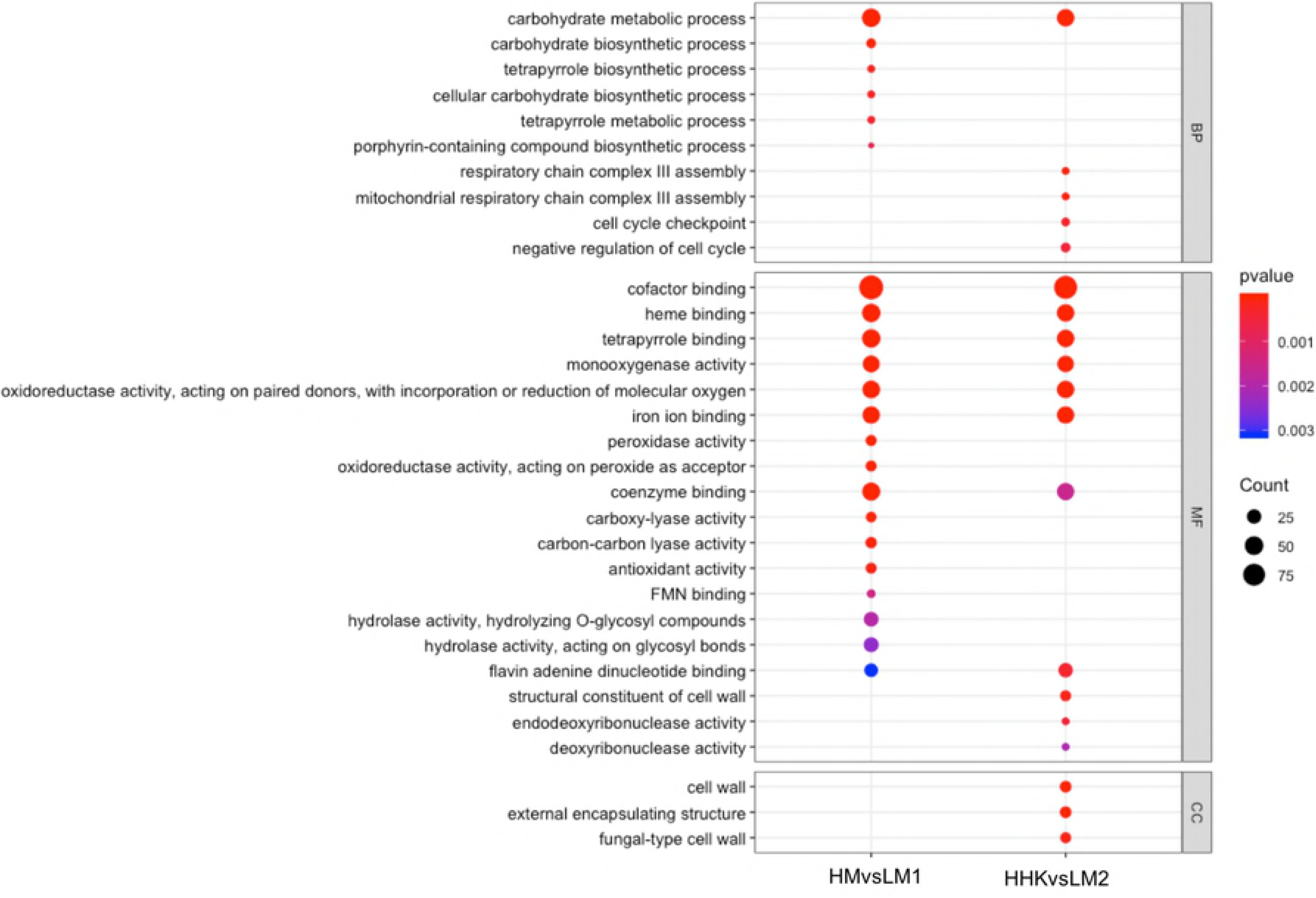
Mycelium radial growth rate of *C. cinerea* in LiCl treated samples and control. Control: treated with water; the concentration of LiCl: 0.25 μM, 0.5 μM, 1 μM, 3 μM and 5 μM. Vertical bars represent standard deviation of the mean (n=3).

### Effect of LiCl on fruiting body initiation

To induce fruiting body formation, plates were transferred to an incubator with 12/12 hrs light/dark cycle at 28°C. After 3-4 days of incubation, initials were observed on the control plates and hyphal knots on plates with 0.25 μM LiCl. By contrast, plates with other concentrations of LiCl were still in the mycelium stage. After 5-6 days, young fruiting bodies were observed on the control plates and primary primordium on the 0.25 μM LiCl plates (Fig. 2). Initials and hyphal knots were observed on the 0.5 μM and 1 μM LiCl plates, respectively. The 3 μM and 5 μM LiCl plates were still in the mycelium stage. Moreover, the mycelial growth in the 5 μM LiCl plates was slightly faster than that in the control plates and the mycelium stopped growth by reaching to the edge of plates. After one week of incubation, the fruiting bodies in the control plates and the plates treated with 0.25 μM, 0.5 μM and 1 μM LiCl autolyzed consecutively whereas the 3 μM and 5 μM LiCl plates remained in the mycelium stage with no fruiting body formation.

**Fig 2.**
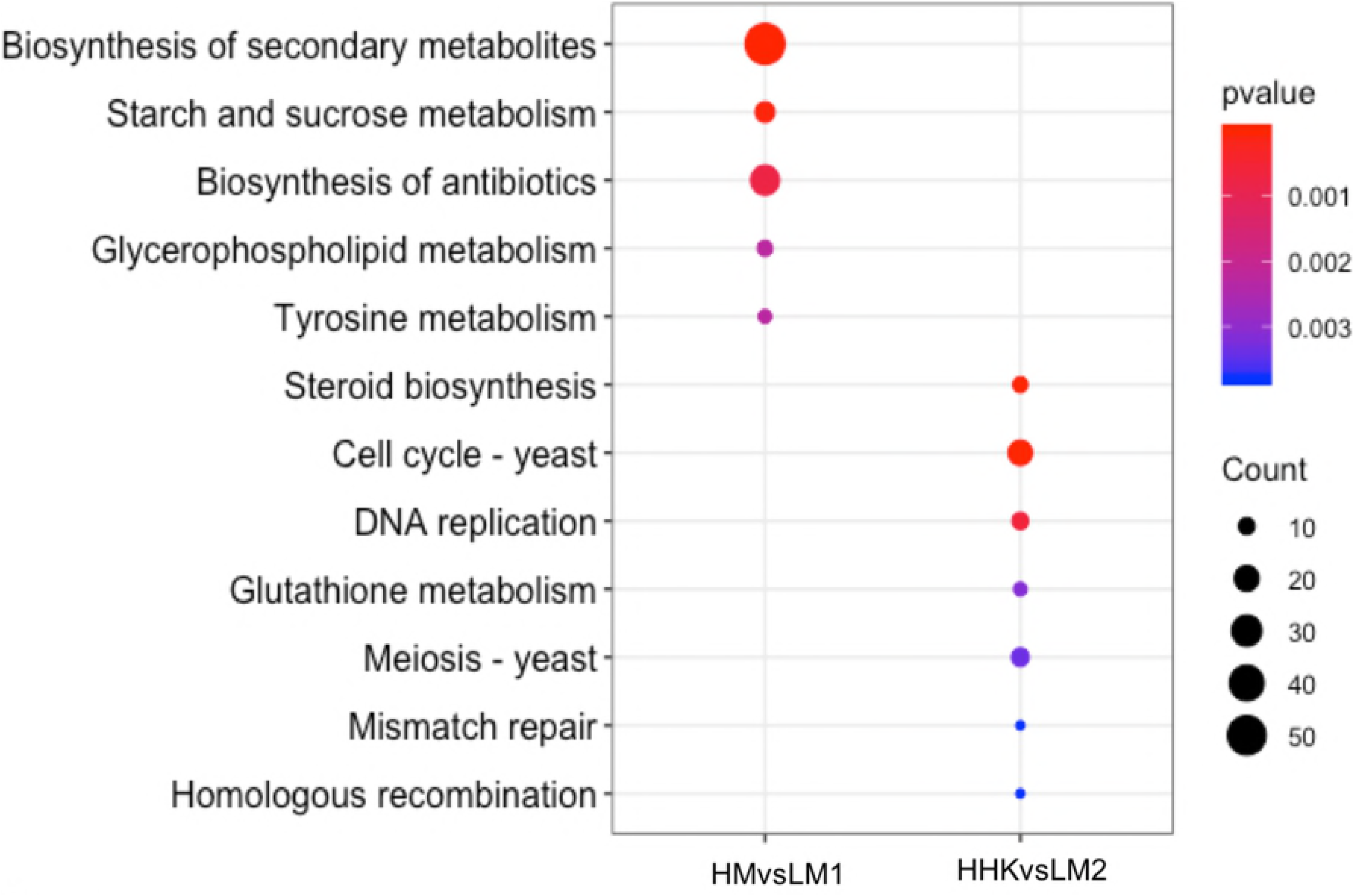
Effect of different concentrations of LiCl in *C. cinerea* fruiting body development. Primordium was observed on the YMG plates treated with 0.25 μM LiCl after 5 days of incubation. Initials and hyphal knots were observed on the YMG plates treated with 0.5 μM and 1 μM LiCl, respectively. YMG plates treated with 3 μM and 5 μM LiCl remained in the mycelium stage, and mycelium treated with 5 μM LiCl stopped reaching the edge of plates. Young fruiting bodies were observed on the control YMG plates treated with water.

### Sequencing of the ***C. cinerea*** transcriptome

To study the transcriptional response of *C. cinerea* to GSK3 inhibition, transcriptomes of *C. cinerea* treated with water and 3 μM LiCl in the mycelium and hyphal knots stages were sequenced on the Illumina HiSeq 2000 platform. RNA-Seq resulted in 2.6-3.8×10^7^ reads and 3.9-5.8×10^9^ bases. Subsequent quality filtering removed 0.02% of the reads that were deemed unclear. About 94% of the reads were mapped to the reference genome from JGI, resulting in 91240 transcripts and 14128 unigenes. All unigenes were annotated to GO and KOG, comprising 2150 functional GO terms and 25 KOG classes.

### Transcriptional profiling of the effect of LiCl on mycelial growth enhancement

Of the 14128 unigenes expressed in the mycelium treated with water (HM) and treated with 3 μM LiCl (LM1), 1210 were significantly differentially expressed (P<0.05, |log2 fold change|>1), including 531 upregulated and 679 downregulated unigenes in LM1(S1 Dataset). GO annotation showed that both the upregulated and downregulated DEGs had a similar distribution in the molecular function (~70%), biological process (~23%) and cellular component (7%) categories (S1 Fig.).

The DEGs were also annotated into 667 KOG terms, including 261 upregulated and 406 downregulated KOG terms. Almost half of the upregulated DEGs were annotated to metabolism, 22% to cellular processes and signaling, and 14% to information storage and processing. Downregulated DEGs had similar number of genes (about 33%) annotated to metabolism and cellular processes and signaling, with 9% annotated to information storage and processing (Fig 3).

**Fig. 3.**
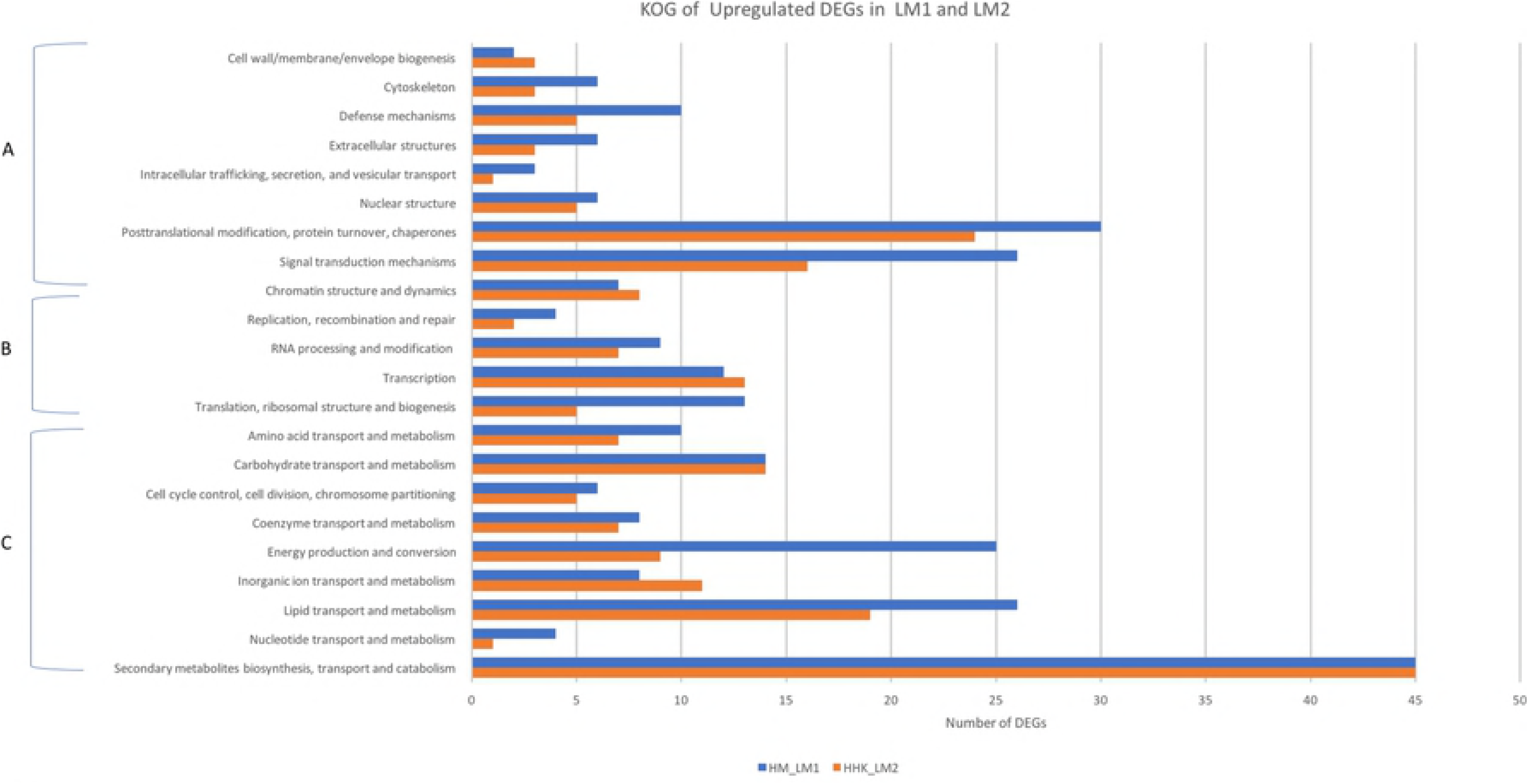
KOG annotation of DEGs. The distribution of KOG annotations of (A) upregulated and (B) downregulated DEGs in HM vs LM1, HM: *C. cinerea* mycelium treated with water, LM1: *C. cinerea* mycelium treated with 3 μM LiCl; and (C) upregulated and (D) downregulated DEGs in HHK vs LM2, HHK: hyphal knots of the water treated samples, LM2: consistent mycelium of 3 μM LiCl-treated samples.

GO functional enrichment analysis resulted in two categories, biological process and molecular function, with P-value ≤0.05 (Fig 4). Most of the DEGs between HM and LM1 were enriched in molecular function, including “cofactor binding”, “heme binding”, “tetrapyrrole binding”, and “coenzyme binding”. Besides, six DEGs were enriched in biological process, including “carbohydrate metabolic process”, “carbohydrate biosynthetic process”, “tetrapyrrole biosynthetic process”, “cellular carbohydrate biosynthetic process”, “tetrapyrrole metabolic process”, and “porphyrin-containing compound biosynthetic process”. No DEGs were significantly enriched in the cellular component category, indicating that cellular component has no significant effect on the transcriptional response of *C. cinerea* mycelium to LiCl treatment.

**Fig.4.**
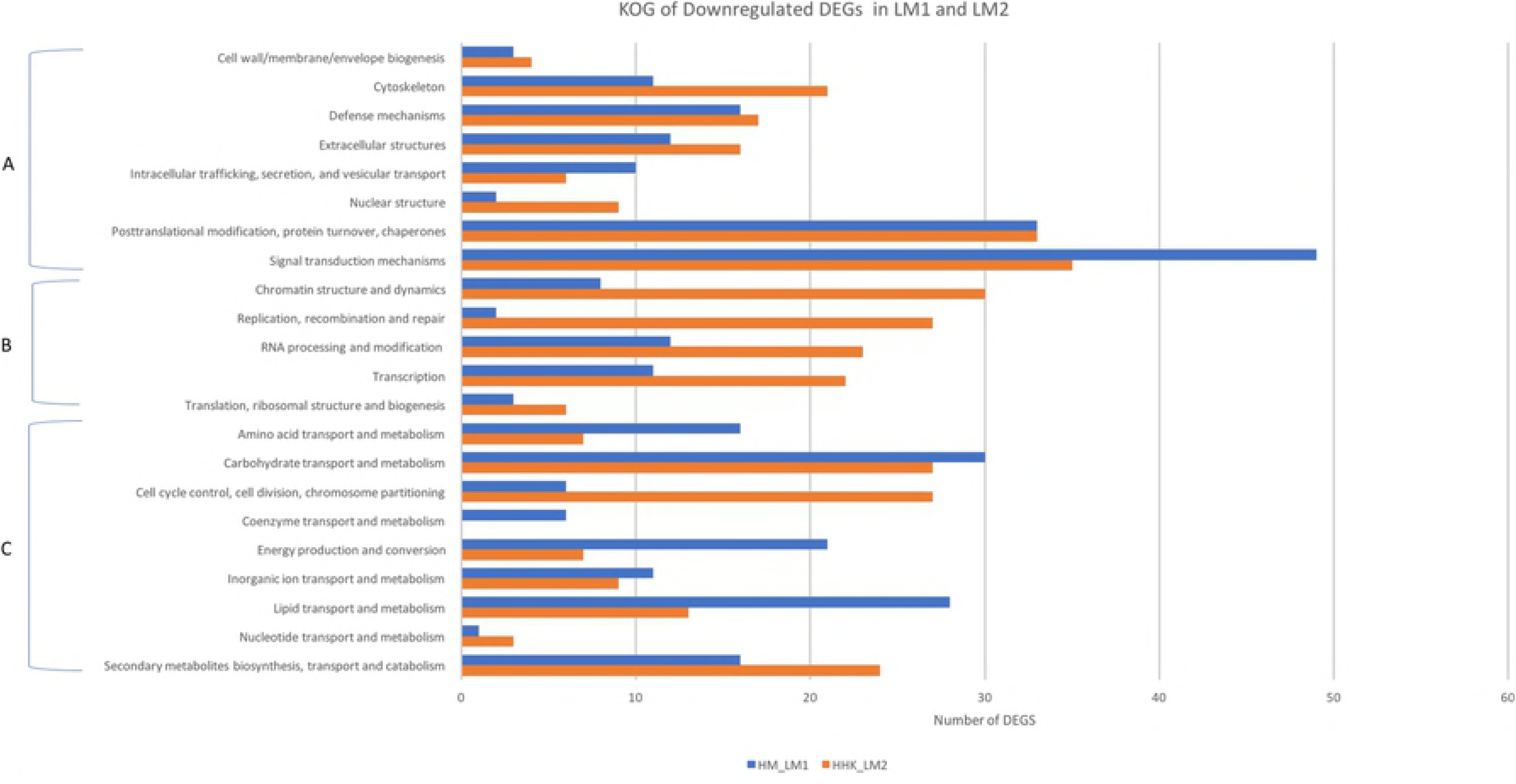
GO functional enrichment analysis. The size of the dots indicates the number of DEGs involved in the pathway and the color represents the P-value. HM: *C. cinerea* mycelium treated with water; LM1: *C. cinerea* mycelium treated with 3 μM LiCl; HHK: hyphal knots of the water treated samples; LM2: consistent mycelium of 3 μM LiCl-treated samples.

KEGG pathway enrichment analysis resulted in five pathways statistically enriched in the LiCl-treated samples with P-value ≤0.05, including “biosynthesis of secondary metabolites”, “biosynthesis of antibiotics”, “starch and sucrose metabolism”, “glycerophospholipid metabolism”, and “tyrosine metabolism” (Fig 5). This result indicates that the enhanced mycelial growth of *C. cinerea* treated with 3 μM LiCl was mainly associated with metabolism pathways, especially those in secondary metabolism.

**Fig 5.**
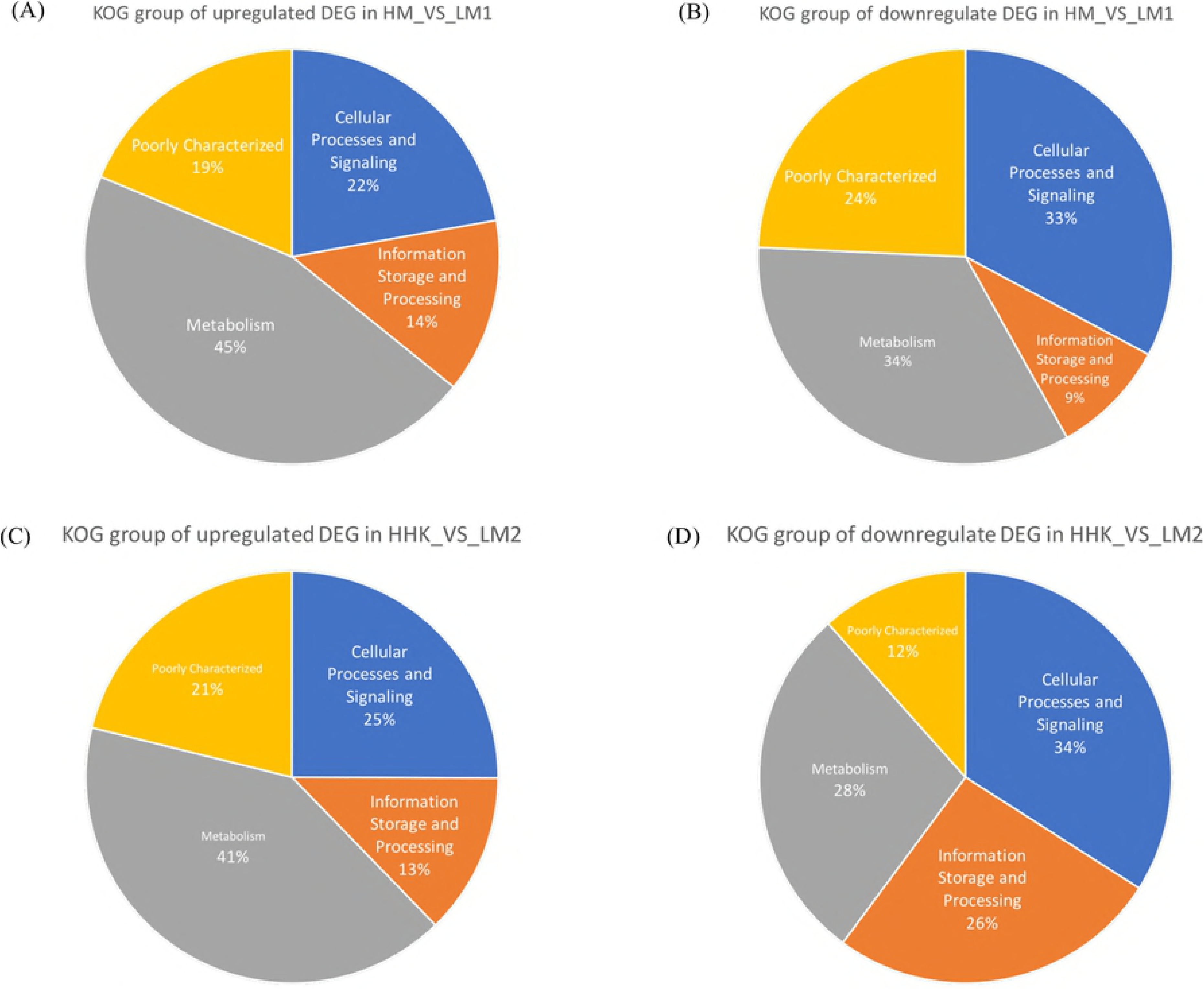
KEGG pathway enrichment analysis. The size of the dots indicates the number of DEGs involved in the pathway and the color represents the P-value. HM: *C. cinerea* mycelium treated with water; LM1: *C. cinerea* mycelium treated with 3 μM LiCl; HHK:

### Transcriptional profiling of the effect of LiCl on fruiting body initiation repression

A total of 1402 unigenes were significantly differentially expressed between hyphal knots of the water treated samples (HHK) and consistent mycelium of LiCl-treated samples (LM2) (P<0.05, |log2 fold change|>1), of which 664 were upregulated and 738 were downregulated in LM2 (S2 Dataset). GO annotation showed that both the upregulated and downregulated DEGs had a similar distribution in the molecular function (71-75%), biological process (17~22%) and cellular component (7~8%) categories (S1 Fig.).

The DEGs were annotated into 769 KOG terms, 355 of which were upregulated and 414 were downregulated in LM2. Forty-one percent of the upregulated DEGs belonged to metabolism, followed by 25% to cellular processes and signaling, and 13% to information storage and processing (Fig 3). For the downregulated DEGs, 34% were involved in cellular process and signaling, 28% in metabolism and 26% in information storage and processing. The number of upregulated DEGs in LM2 annotated to KOG classes were less than that in LM1, however, the number of downregulated DEGs in LM2 annotated to KOG classes were greater than that in LM1 (Fig 6 &7).

**Fig 6.**
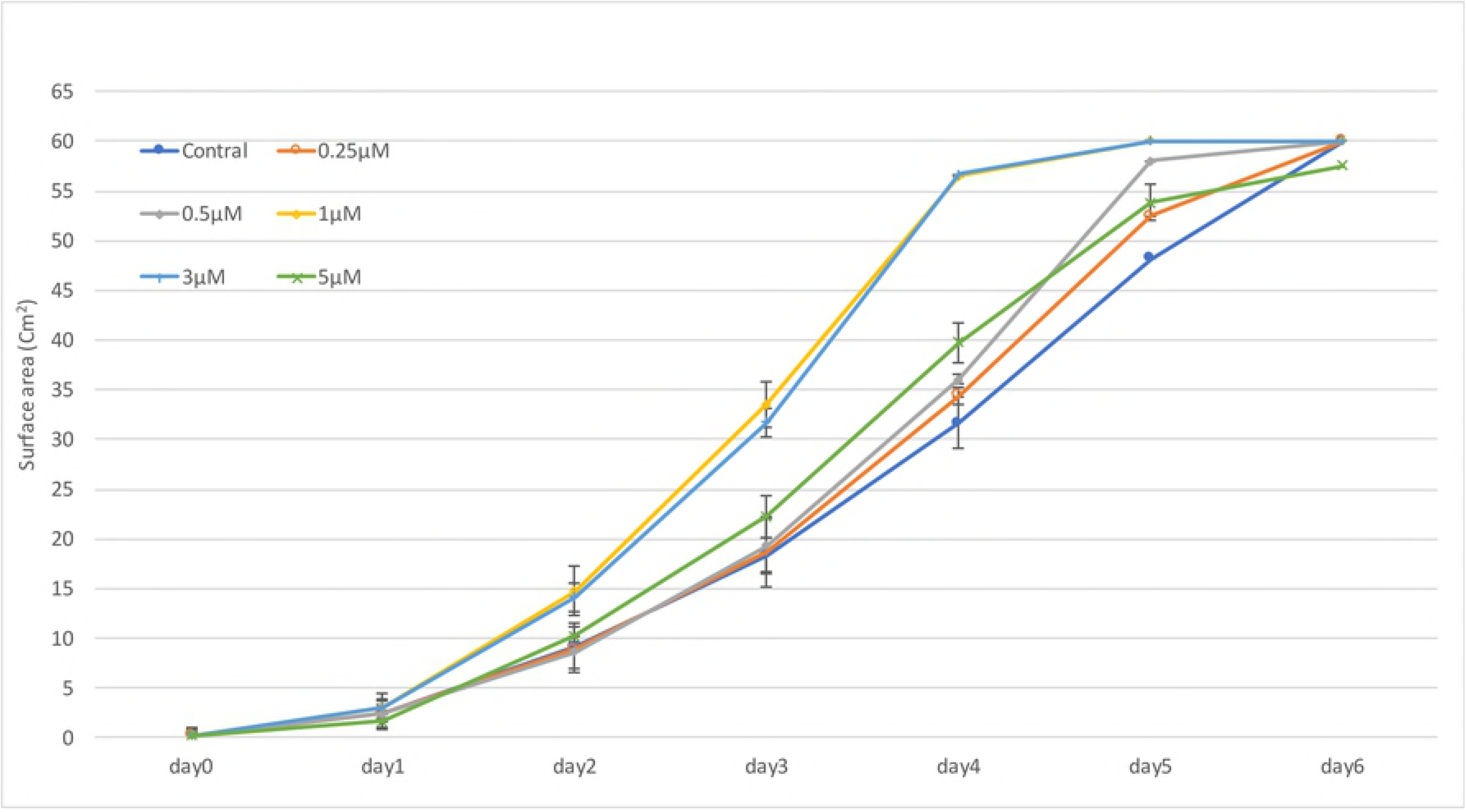
KOG classification of upregulated DEGs in the LiCl-treated samples. A: Cellular processes and signaling; B: Information storage and processing; C: Metabolism. HM: *C. cinerea* mycelium treated with water; LM1: *C. cinerea* mycelium treated with 3 μM LiCl;

**Fig 7.**
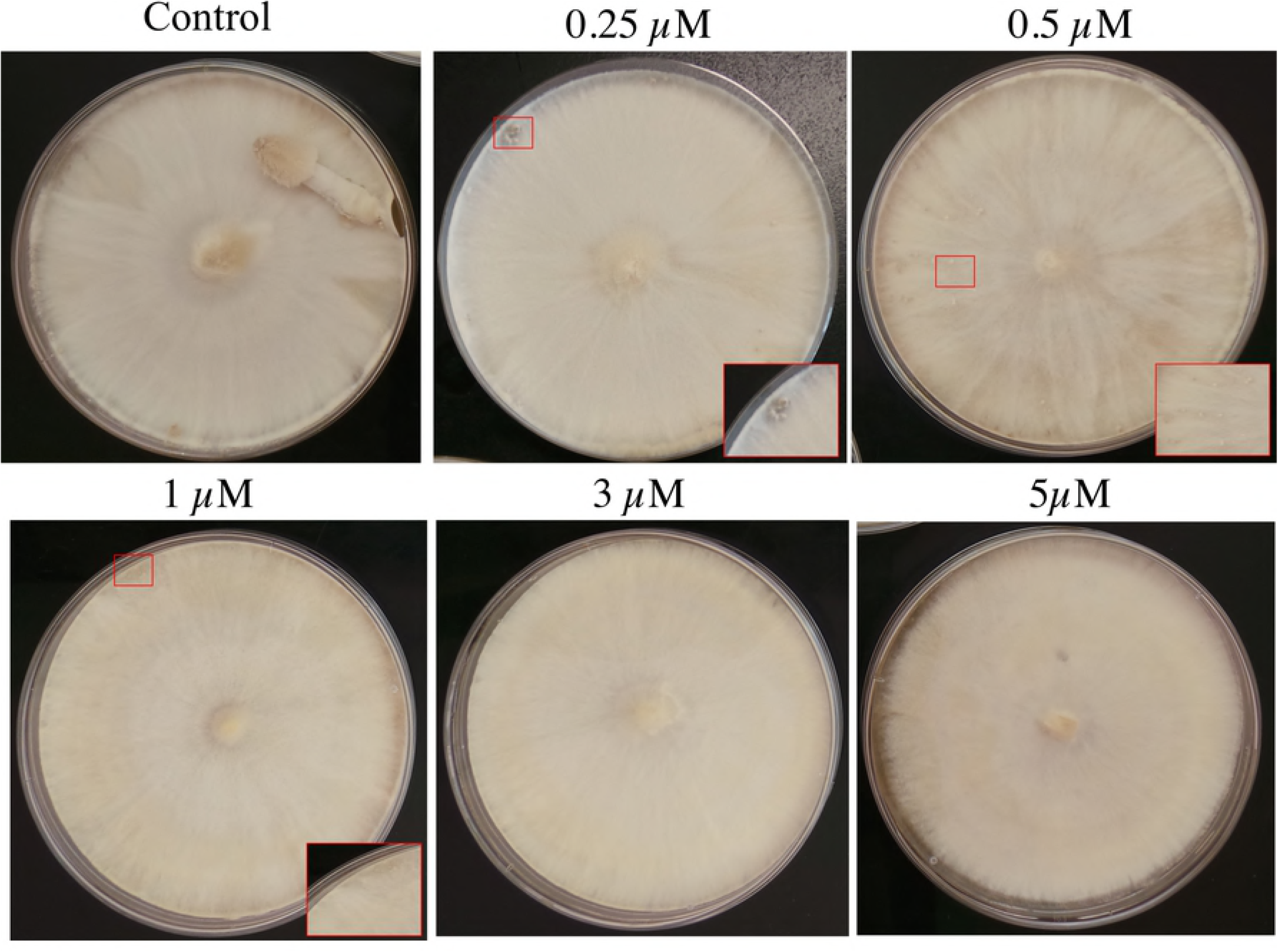
KOG classification of downregulated DEGs in the LiCl-treated samples. Cellular processes and signaling; B: Information storage and processing; C: Metabolism; HM: *C. cinerea* mycelium treated with water; LM1: *C. cinerea* mycelium treated with 3 μM LiCl; HHK: hyphal knots of the water treated samples; LM2: consistent mycelium of 3 μM LiCl-treated samples.

GO functional enrichment analysis resulted in 11 statistically enriched GO terms from the molecular function category, including “cofactor binding”, “iron ion binding”, “heme binding”, and “structural constituent of cell wall”. Five enriched GO terms were from the biological process category, namely “carbohydrate metabolic process”, “respiratory chain complex III assembly”, “mitochondrial respiratory chain complex III assembly”, “cell cycle checkpoint”, and “negative regulation of cell cycle”. Moreover, three enriched GO terms were from the cellular component category, namely “cell wall”, “external encapsulating structure” and “fungal-type cell wall”. Eight GO terms from the molecular function category were enriched in both time points, namely “cofactor binding”, “heme binding”, “tetrapyrrole binding”, “iron ion binding”, “coenzyme binding, “Flavin adenine dinucleotide binding”, “monooxygenase activity”, and “oxidoreductase activity”, indicating their continuous involvement in fruiting body initiation in *C. cinerea* (Fig 4.).

KEGG pathway enrichment analysis of DEGs between HHK and LM2 resulted in seven statistically enriched pathways in the LiCl-treated samples (Fig 5). These included “steroid biosynthesis” and “glutathione metabolism” from the metabolism pathways category and “Cell cycle-yeast”, “DNA replication”, “Meiosis-yeast”, “Mismatch repair”, and “Homologous recombination” from the cellular process category. All DEGs involved in the cellular process pathways were downregulated (S5 Table), indicating that LiCl suppressed the expression of these DEGs and the inactivation of GSK3 may affect their expression in fruiting body induction.

### Validation of the RNA-Seq results by qRT-PCR

Eight genes involved in the enrichment analysis at the mycelium and hyphal knots stages were selected to validate the results of transcriptional profiling. The expression patterns of genes revealed by qRT-PCR were consistent with those in the RNA-Seq analysis, indicating that the results of transcriptional profiling analysis are reliable (S5 Table).

## Discussion

To reveal the involvement of GSK3 in the fruiting body development of *C. cinerea*, we treated *C. cinerea* mycelium with a GSK3 inhibitor, Lithium chloride (LiCl). The result showed that mycelium treated with LiCl grew faster than that of the control. KEGG enrichment analysis showed that the enriched DEGs in LM1 in metabolism pathways. Two enzymes were upregulated and six were downregulated in the “starch and sucrose metabolism pathway” (S2 Fig.). In the glycogen synthesis process, UDP-glucose is the substrate of glycogen synthase [31]. The downregulated expression of glycogen synthase detected here may then allow more UDP-glucose to be used in the synthesis of beta (1,3)-D-glucan, a component of the cell wall, for mycelium growth [23]. Besides, the upregulation of glucoamylase and downregulation of alpha-amylase allow breakdown of glycogen into glucose for mycelium growth instead of dextrin. Moreover, the upregulation of beta-glucosidase breaks down beta-D-glucosides or cellobiose into glucose, thereby allowing rapid cell division and accumulation of cell mass [24]. Moreover, galectin regulates cell growth and extracellular cell adhesion [25]. The expression of galectin in mycelium is low and it is upregulated in nutrient depletion for hyphal aggregation in previous studies [26, 27]. In this study, galectin-1 (CC1G_05003) and galectin-2 (CC1G_05005) (S1Dataset) were upregulated in the LiCl-treated samples, it implies that the phosphorylation of GSK3 suppresses the expression of galectin in mycelium under excessive nutrient.

The time required for fruiting body formation increased when the concentration of LiCl increased. Upon 3 μM LiCl, *C. cinerea* fruiting body formation was inhibited even under favorable environmental conditions. When mycelium transits to hyphal knots, it required expression of several genes for the construction of new membranes. In a previous transcriptome study, genes involved in the phospholipid biosynthesis process were upregulated in the fruiting body initiation of *C. cinerea* [3]. In this study, genes of the phospholipid biosynthesis process, including phosphatidylserine decarboxylase (CC1G_06249, CC1G_09238, CC1G_09240, and CC1G_09237), phosphatide cytidylyltransferase (CC1G_06860), inositol-3-phosphate synthase (CC1G_06001), and a hypothetical protein (CC1G_09235) (S6 Table) were downregulated in LM2 treated with 3 μM LiCl compared with the control samples that formed hyphal knots. Most of these genes participate in glycerophospholipid metabolism. Therefore, we suggest that GSK3 activity affects glycerophospholipid metabolism for the transition from mycelium to hyphal knots. Besides, fruiting body induction requires some important chromosome remodeling genes [2,28,29]. Those genes, however, were downregulated in this study (S6 Table), indicating that the activity of GSK3 might associate with the expression of chromosome remodeling genes for cell division, which results in hyphal knots formation [30].

KEGG pathway enrichment analysis showed that DEGs involved in “cell cycle-yeast”, “meiosis-yeast”, “DNA replication”, “mismatch repair”, and “homologous recombination” in the cellular process category were downregulated in LM2 compared to the controls, showing that *C. cinerea* treated with LiCl remained in the mycelium stage as a result of the downregulation of genes involved in cell cycle and meiosis. Phosphorylation of a yeast GSK3 homolog has been found to promote vegetative growth in cell cycle and meiotic development pathway [32,33]. In *Fusarium. graminearum*, the signal GSK3 homolog is suggested to promote the transcription of numerous early meiosis-specific genes as the Δfgk3 mutant failed in sexual development processes [9,34]. In this study, we suggest that the phosphorylation of GSK3 is essential for cellular processes during fruiting body formation as the inhibition of GSK3 activity by LiCl in *C. cinerea* suppressed fruiting body formation.

KOG analysis showed a similar distribution of upregulated DEGs involved in four KOG categories at both time points, indicating that gene upregulation does not play a significant role in the inhibition of fruiting body initiation via GSK3 inactivation. On the other hand, the increased contribution of downregulated genes in information storage and processing at the second time point indicates that the inhibition of fruiting body initiation is via downregulation of relevant genes. The contribution of DEGs involved in cellular processes and signaling was similar between the two time points, indicating that the activity of GSK3 inhibited by LiCl constantly downregulated genes involved in cellular processes and signaling despite changes in environmental conditions. At first, the mycelia were incubated in nutrient-rich plates at a high temperature under total darkness. After the mycelia reached the edge of plates, the plates were then transferred to a nutrient-reduced, low temperature environment under light/dark cycle. Due to the environmental changes, genes involved in cellular processes and signaling responded to initiate fruiting body formation. The inhibition of GSK3 suppressed the expression of genes involved in cellular processes and signaling even when the environmental conditions were changed to a favorable condition for fruiting body initiation. Therefore, we suggest that the activity of GSK3 is required for fruiting body initiation. However, how environmental factors such as light, temperature and nutrient interact with GSK3 to stimulate the global gene expression changes for fruiting body induction requires further investigation.

Fruiting body formation in basidiomycetes is influenced by individual or multiple environmental factors, such as light, temperature and nutrients. In this study, *C. cinerea* was first grown at 37°C under total darkness with rich nutrient and then at 28°C in a light/dark cycle after the mycelium reached the edge of plates. Light, cold shock and nutrient diminution trigger or promote fruiting as well as the expression of stress response genes [1, 27, 35, 36]. Several studies have found that the expression of stress response genes to environmental conditions are upregulated during fruiting body formation [1,3,37]. However, in this study, the expression of nutrient response genes such as PKA protein kinase (CC1G_01089) were significantly deregulated and adenylate cyclase (CC1G_02340) were slightly downregulated in LM2 treated with LiCl compared with the controls (S2 Dataset). Both participant in cAMP-PKA pathway and involve in sensing the nutrient level during development transition [37]. As GSK3 homologs in yeast and *Neurospora crassa* take a response in nutrient signal and light signal transmission [11,12,13], we propose that the activity of GSK3 may be an important channel between extracellular signals to respective response gene expression related to fruiting body formation.

## Conclusion

To the best of our knowledge, this study is the first transcriptome profiling of the role of GSK3 in *C. cinerea* fruiting formation. We found that the inhibition of GSK3 activity by the addition of LiCl at the mycelium stage enhanced the mycelial growth. This phenotype could be explained by the enriched metabolism pathways, such as “biosynthesis of secondary metabolite” and “biosynthesis of antibiotic”, in which DEGs were upregulated to release glucose for energy. Moreover, DEGs involved in cellular process pathways, including “cell cycle-yeast” and “meiosis-yeast”, were identified in the *C. cinerea* fruiting body formation suppressed by LiCl under favorable environmental conditions. Our data suggest that the activity of GSK3 is essential for fruiting body formation by affecting the expression of fruiting body induction genes and genes in cellular processes. Further studies on the phosphorylation of GSK3 with external environmental signals may facilitate the understanding of how environmental factors induce fruiting body development.

## Acknowledgement

We thank Dr. Hajimme Muraguchi for sharing *C. cinerea* strains used in this study.

## Supporting information

S1 Dataset. Differentially expressed genes in LM1 against HM, HM: *C. cinerea* mycelium treated with water; LM1: *C. cinerea* mycelium treated with 3 μM LiCl.

S2 Dataset. Differentially expressed genes in LM2 against HHK, HHK: hyphal knots of the water treated samples; LM2: consistent mycelium of 3 μM LiCl-treated

S1 Table. Primers designed for validation of gene expression profile in *C. cinerea* transcriptome data.

S2 Table. Unigenes used for validation of gene expression profile in *C. cinerea* transcriptome data.

S3 Table. KEGG enrichment between different samples (p-value ≤ 0.05).

S4 Table. GO enrichment between different samples (p-value ≤ 0.05).

S5 Table. DEGs involved in KEGG enriched pathways between HHK and LM2.

S6 Table The downregulated expression gene of fruiting body induction based on RNA-seq analysis.

**S1 Fig. Go annotation of DEGs.** The distribution of GO annotations of (A) upregulated and (B) downregulated DEGs in HM vs LM1, and (C) upregulated and (D) downregulated DEGs in HHK vs LM2.

**S2 Fig. One of enriched KEGG pathways in HM vs LM1 transcriptome analysis.** Starch and sucrose metabolism pathway is one of the enriched metabolism pathways in samples treated with 3 μM LiCl. Red boxes indicate upregulated DEGs and green boxes indicate downregulated DEGs in HM vs LM1. Gray boxes indicate non-DEGs.

